# Bupropion inhibits serotonin type 3AB heteromeric channels at a physiologically relevant concentration

**DOI:** 10.1101/709881

**Authors:** Antonia G. Stuebler, Michaela Jansen

## Abstract

Bupropion, a FDA-approved antidepressant and smoking cessation aid, blocks dopamine and norepinephrine reuptake transporters and non-competitively inhibits nicotinic acetylcholine (nACh) and serotonin type 3A (5-HT_3_) receptors. 5-HT_3_ receptors are pentameric ligand-gated ion channels that regulate synaptic activity in the central and peripheral nervous system pre- and postsynaptically. In the present study, we examined and compared the effect of bupropion and its active metabolite hydroxybupropion on homomeric 5-HT_3A_ and heteromeric mouse 5-HT_3AB_ receptors expressed in *Xenopus laevis* oocytes using two-electrode voltage clamp experiments. Co-application of bupropion or hydroxybupropion with 5-HT dose-dependently inhibited 5-HT-induced currents in 5-HT_3AB_Rs (IC_50_ = 866 μM and 505 μM, respectively) but potentiated 5-HT-induced currents at low (30-50 μM) concentrations. The corresponding IC_50_s for bupropion and hydroxybupropion with 5-HT_3A_R were 10- and 5-fold lower, respectively (87 μM and 113 μM), and no potentiation was observed. The inhibition of 5-HT_3A_R and 5-HT_3AB_R was non-use dependent and voltage-independent, indicating bupropion is not an open channel blocker. The inhibition by bupropion was reversible and time-dependent. Of note, pre-incubation with a low concentration of bupropion that mimics therapeutic drug conditions significantly inhibited 5-HT induced currents in 5-HT_3A_ and even more so 5-HT_3AB_ receptors. In summary, our results indicate that bupropion inhibits 5-HT_3AB_R, as well as homomeric receptors, and that this inhibition takes place at clinically-relevant concentrations. Inhibition of 5-HT_3_ receptors by bupropion may contribute to its desired and/or undesired clinical effects.

**Significance Statement:** 5-HT_3AB_ receptors are found in brain areas involved in mood regulation. Clinical studies indicate that antagonizing these receptors was successful in treating mood and anxiety disorders. Some currently clinically available antidepressants and antipsychotics act as antagonists of 5-HT_3_ receptors. Previously, bupropion was shown to be an antagonist at homopentameric 5-HT_3A_ receptors. The present work provides novel insights into the pharmacological effects bupropion exerts on heteromeric 5-HT_3AB_ receptors. The results advance the knowledge on the clinical effect of bupropion as an antidepressant.

## Introduction

The 5-hydroxytryptamine-3 (5-HT_3_) receptor is an ionotropic receptor and a member of the Cys-loop family of pentameric ligand-gated ion channels (pLGICs) and thereby differs from G-protein-coupled serotonin receptors (5-HT_1_ to 5-HT_7_) (Hannon, 2008; Hoyer, 2002; Thompson, 2007). 5-HT_3_R is similar in structure and function to the other members of the pLGICs, including the cation-selective nicotinic acetylcholine receptor (nAChR) and the anion-selective γ-amino butyric acid (GABA_A_) and glycine receptors. Malfunction in these receptors has been linked to several neurological disorders (Hogg, 2003; Lemoine, 2012). Together, they are responsible for fast neurotransmission in the central and peripheral nervous system (Thompson, 2013) and are involved in virtually all brain functions (Hassaine, 2014).

To date, five different 5-HT_3_ subunits have been identified (5-HT_3A_ – 5-HT_3E_). The first subunit to be cloned, 5-HT_3A_ (Maricq, 1991), is the only subunit that can form functional homo-oligomeric receptors on the cell membrane when expressed in *Xenopus* oocytes or cell lines (Hussy, 1994). Introduction of the 5-HT_3B_ subunit, to form functional heteromers, altered the properties of the homo-oligomer that more closely resembled the functional responses observed in native tissues (Davies, 1999; Dubin, 1999; Hussy, 1994; Maricq, 1991). When compared to 5-HT_3A_R, 5-HT_3AB_R differs in agonist concentration-response curves, shows increased single-channel conductance and desensitization, and an altered current-voltage relationship (Davies, 1999; Dubin, 1999; Kelley, 2003b; Thompson, 2013).

The 5-HT_3_R is widely distributed in the central and peripheral nervous systems, and on extraneuronal cells, such as monocytes, chondrocytes, T-cells, and synovial tissue (Fiebich, 2004). In the periphery, 5-HT_3_Rs are found in the autonomic, sensory and enteric nervous systems (Faerber, 2007) where they are involved in regulating gastrointestinal (GI) functions, such as motility, emesis, visceral perception and secretion (Karnovsky, 2003; Lummis, 2012; Niesler, 2011; Niesler, 2003; Niesler, 2007). The highest density of 5-HT_3_Rs in the CNS is in the hindbrain, particularly the dorsal vagal complex involved in the vomiting reflex, and in limbic structures, notably the amygdala, hippocampus, nucleus accumbens and striatum (Barnes, 1989; Hannon, 2008; Jones, 1992; Miyake, 1995). Significant 5-HT_3B_ expression was identified in the human brain with high levels in the amygdala, hippocampus, and the nucleus caudate (Davies, 1999; Dubin, 1999; Niesler, 2003; Tzvetkov, 2007). A high amount of 5-HT_3_ is found on pre-synaptic nerve fibers (Miquel, 2002; Nayak, 2000), through which it can modulate the release of other neurotransmitters, such as dopamine, cholecystokinin, GABA, substance P and acetylcholine (Chameau, 2006; Faerber, 2007). Owing to its involvement in many brain functions, the 5-HT_3_R represents an attractive therapeutic target.

5-HT_3_R antagonists are used to effectively treat patients experiencing irritable bowel syndrome and chemotherapy-/radiotherapy-induced and postoperative nausea and vomiting (Thompson, 2007). Some antidepressants (Choi, 2003; Eisensamer, 2003) and antipsychotic drugs (Rammes, 2004) also antagonize 5-HT_3_R, which, together with other preclinical and clinical studies, suggest the relevance of 5-HT_3_R antagonism for treating psychiatric disorders (Bétry, 2011). We recently discovered that bupropion, another antidepressant, antagonizes 5-HT_3A_R (Pandhare, 2017).

Bupropion was first approved as an ‘atypical’ antidepressant over 30 years ago, and today is one of the most commonly prescribed antidepressants. Despite its recognized clinical efficacy for both depression and smoking cessation, a comprehensive picture of how bupropion modulates neurotransmission is still emerging. Bupropion’s therapeutic effect is thought to be mediated by the blocked reuptake of dopamine (DA) and norepinephrine (NE) (Stahl, 2004) and the non-competitive inhibition of neuronal and muscular AChRs (Alkondon, 2005; Arias, 2009b; Fryer, 1999; Slemmer, 2000).Most recently, the discovery that bupropion also non-competitively inhibits 5-HT_3A_R (Pandhare, 2017) raises the question of whether this inhibition takes place at physiologically relevant concentrations and if bupropion inhibits the heteromeric members of the 5-HT_3_ family. Therefore, we investigated the effect of bupropion and its major metabolite, hydroxybupropion, on the function of heteromeric 5-HT_3AB_Rs as compared to the homomeric receptor in *Xenopus* oocytes. Here, we demonstrate that 5-HT_3AB_R, like 5-HT_3A_R, is dose-dependently inhibited by bupropion and its metabolite. This inhibition is voltage-independent, non-use dependent (i.e. affected by pre-incubation), and occurs significantly at physiologically relevant concentrations.

## Materials and Methods

### Materials

Protease inhibitor cocktail III (Research Products International, Mt. Prospect, IL); bupropion hydro-chloride and hydroxybupropion (Toronto Research Chemicals, Inc. North York, Canada); serotonin (5-HT; serotonin creatinine sulfate monohydrate) from Acros Organics (New Jersey, NJ). Stock of serotonin (2 mM) and bupropion (100 mM) were prepared in distilled water. Hydroxubupropion (100 mM) was dissolved in dimethyl sulfoxide (DMSO). All solutions were made in OR-2 buffer immediately before conducting experiments.

### Molecular Biology

Complementary DNA encoding the mouse 5-HT_3A_R, containing a V5-tag (GKPIPNPLLGLDSTQ) close to the N-terminus (Jansen, 2008), and 5-HT_3AB_R in the pGEMHE vector were used for oocyte expression (Reeves, 2001). Plasmids were linearized with the *Nhe*I restriction enzyme and *in vitro* transcribed with the T7 RNA polymerase kit (mMESSAGE mMACHINE^®^ T7 Kit; Applied Biosystems/Ambion, Austin, TX). Capped cRNA was purified with the MEGAclear™ Kit (Applied Biosystems/Ambion, Austin, TX), and precipitated using 5 M ammonium acetate. cRNA dissolved in nuclease-free water was stored at −80 °C.

### *Xenopus laevis* oocyte preparation

Oocytes were isolated, enzymatically defolliculated, and stored as previously described (Goyal, 2011). *Xenopus laevis* frogs were handled and maintained following procedures approved by the local animal welfare committee (Institutional Animal Care and Use Committee, IACUC #: 08014, PHS Assurance # A 3056-01). In brief, the isolated oocytes were incubated with collagenase (Collagenase from *Clostridium histolyticum* Type IA, Sigma-Aldrich) for one hour in OR-2 (115 mM NaCl, 2.5 mM KCl, 1.8 mM MgCl2, 10 mM HEPES, pH 7.5), followed by extensive washing with OR-2. Oocytes were then rinsed three times with OR-2 + 2 mM Ca^2+^ for 45min each and maintained in standard oocyte saline medium (SOS: 100 mM NaCl, 2 mM KCl, 1 mM MgCl_2_, 1.8 mM CaCl_2_, 5 mM HEPES, pH 7.5, supplemented with 1% antibiotic-antimycotic, 5% horse serum) for up to 7 days at 16 °C. Oocytes were microinjected with 10 ng of *in vitro* synthesized mRNA (0.2 ng/μL) using an automatic oocyte injector (Nanoject II™; Drummond Scientific Co., Broomall, PA) up to 48 hours after isolation. For optimal expression of the heteromeric 5-HT_3AB_R, the A and B subunits were co-injected in a 1:3 ratio. This ratio has proven to be optimal for 5-HT_3AB_R expression, as a lower ratio results in 5-HT_3A_R mimicked current response and higher ratio would impact overall receptor expression (Corradi, 2015; Thompson, 2013).

### Electrophysiology

Two-electrode voltage clamp recordings were performed and analyzed using a TEV-200A amplifier (Dagan Instruments, Minneapolis, MN), a Digidata 1440A data interface (Molecular Devices, Sunnyvale, CA), MiniDigi 1B (Molecular Devices) and pClamp 10.7 software (Molecular Devices). Recordings were conducted 1 – 4 days after microinjection. All experiments were performed at room temperature (RT, 22–24°C) and at a holding potential of −60 mV, unless otherwise stated. The oocytes were held in a 250 µL chamber and perfused with base solutions (OR-2) using gravity flow at an approximate rate of 5mL/min. Drugs and serotonin were dissolved in the same solution and applied by gravity perfusion. All solutions were applied to a single oocyte until currents reached a steady state before switching solutions. The glass microelectrodes were filled with 3 M KCl and had a resistance of below 2 MΩ. Agonists/antagonists were applied until stable response or desensitization was observed to record maximal current response.

### Oocyte membrane preparation

Oocytes expressing 5-HT_3A_R and 5-HT_3AB_R were homogenized and the membranes were isolated as described before (Pandhare, 2017). The oocytes were homogenized with a 0.5 mL pestle in ice-cold vesicle dialysis buffer (VDB: 100 uL/oocyte: 96 mM NaCl, 0.1 mM EDTA, 10 mM MOPS, 0.02% NaN_3_, pH 7.5), supplemented with protease inhibitor cocktail III [2 uL/mL]) and centrifuged (800 g for 10 min at 4 °C). The supernatant was collected and the membranes containing the receptors were obtained by high-speed centrifugation (39,000*g* for 1 h at 4 °C). Membranes were resuspended in VDB (3 μL/ oocyte) containing PI cocktail and stored at −80 °C.

### Western Blotting

For SDS-electrophoresis, 3 μL of the resuspended membrane pellets containing the receptors were run alongside membrane pellet of uninjected oocyte on 4–15% precast gradient TGX Stain-Free™ gels (Bio-Rad Laboratories) for 34 min at 200 V. Proteins were transferred to PVDF membrane using the Trans-Blot Turbo Transfer System (Bio-Rad). The membranes were blocked with 5% blotting-grade blocker (Bio-Rad) in Tris-buffered Saline with Tween (TTBS) (100 μM Tris pH 7.5, 0.9% NaCl, 1% Tween-20) [25 – 30 mL/membrane]) for one hour up to overnight. The membranes were incubated with the primary V5 HRP-conjugated antibody (1:5,000, V5-HRP antibody, invitrogen) in TTBS (5% blotting-grade blocker) for one hour at RT. After the removal of the primary antibody, the membranes were washed 4x for 5min each with TTBS and one 5-min wash with TBS. Proteins were visualized by chemiluminescent detection (Image Quant) of peroxidase substrate activity (SuperSignal West Femto Maximum Sensitivity Substrate, Thermo Scientific).

### Data Analysis

All electrophysiological data was analyzed with pClamp, Origin (OriginLab Corporation; Northampton, MA) and Prism 6 Software (GraphPad Prism^®^; La Jolla, CA). Data is represented as the mean ± S.E.M, maximal current induced by 5-HT was used as the normalizing standard (100% current response) for other current responses in the same oocyte. Statistical significance was determined with paired or unpaired t-test (in Origin) with a cutoff for significance of 0.05 (*p ≤ 0.05 **p ≤ 0.01 ***p ≤ 0.001). The 5-HT (agonist stimulation – 1a), bupropion, or hydroxybupropion (antagonist inhibition – 1b) concentration dependence on 5-HT_3_ currents was fitted using the variable-slope sigmoidal dose response curve equations:

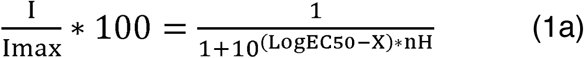

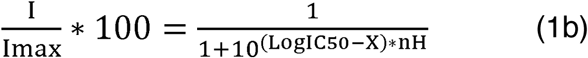

Where I_max_ is the current activated at saturating 5-HT concentration, EC_50_ is the agonist concentration producing 50% of the maximal Imax, IC_50_ is the concentration of antagonist producing 50% inhibition of I_max_, X is the agonist (1a) or antagonist (1b) concentration, and n_H_ is the Hill coefficient. All figures and graphs were made in Origin and Adobe Illustrator CC 2018.

## Results

### Differentiating between 5-HT_3A_R and 5-HT_3AB_R

To evaluate the effect of bupropion and its major metabolite hydroxybupropion (Fig 1) on homomeric and heteromeric 5-HT_3_ receptors, we expressed the mouse 5-HT_3A_R and 5-HT_3AB_R (in a 1:3 A to B ratio (Corradi, 2015; Thompson, 2013)) in *Xenopus* oocytes. First, we substantiated the obvious difference between the two receptor types (Fig 1). The application of the agonist serotonin (5-HT) to *Xenopus* oocytes expressing 5-HT_3A_R (Fig 1A, top trace) or 5-HT_3AB_R (Fig 1A, bottom trace) elicits a rapid inward current with a concentration-dependent amplitude. The currents resulting from a range of 5-HT-concentrations were used to calculate the concentrations that produce a half-maximal response (Fig 1B), yielding EC_50_ values of 0.80 μM (n= 5, Hill slope [n_H_] of 2.53±0.26) for 5-HT_3A_R and 4.30 μM (n = 8; n_H_ = 1.04±0.02) for 5-HT_3AB_R (Table 1). Our EC_50_ value for mouse 5-HT_3A_R and 5-HT_3AB_R are comparable to other values reported previously (Hanna, 2000; Hayrapetyan, 2005; Jansen, 2008; Lochner, 2010; Thompson, 2008). In rodents, the difference between EC_50_ values for 5-HT_3A_ and 5-HT_3AB_ receptors are smaller as compared to the human subunits, nonetheless, the heteromeric receptor shows fast characteristic desensitization kinetics (Fig 1C) and a right-shift in potency (Fig 1B).

**Table 1.** EC_50_ (5-HT) and IC_50_ (bupropion and hydroxybupropion) of *Xenopus laevis* oocytes expressing 5-HT_3A_R and 5-HT_3AB_R

|  | Agonist | pEC<sub>50</sub> (μM,<br>mean±S.E.M.) | EC<sub>50</sub> | n<sub>H</sub><br>(mean±S.E.M.) | n |
| --- | --- | --- | --- | --- | --- |
| 5-HT<sub>3A</sub>R | 5-HT | 6.10±0.02 | 0.80 | 2.53±0.26 | 5 |
| 5-HT<sub>3AB</sub>R |  | 5.38±0.02 | 4.30 | 1.04±0.02 | 8 |
|  | Antagonist | pIC<sub>50</sub> (μM,<br>mean±S.E.M.) | IC<sub>50</sub> | n<sub>H</sub><br>(mean±S.E.M.) | n |
| 5-HT<sub>3A</sub>R | Bupropion | 4.06±0.05 | 87.1 | 1.28±0.07 | 5 |
| 5-HT<sub>3AB</sub>R |  | 3.07±0.04 | 866 | 2.07±0.20 | 7 |
| 5-HT<sub>3A</sub>R | Hydroxybupropion | 3.95±0.04 | 113 | 1.17±0.06 | 5 |
| 5-HT<sub>3AB</sub>R |  | 3.31±0.02 | 505 | 1.57±0.12 | 8 |

**Figure 1.**
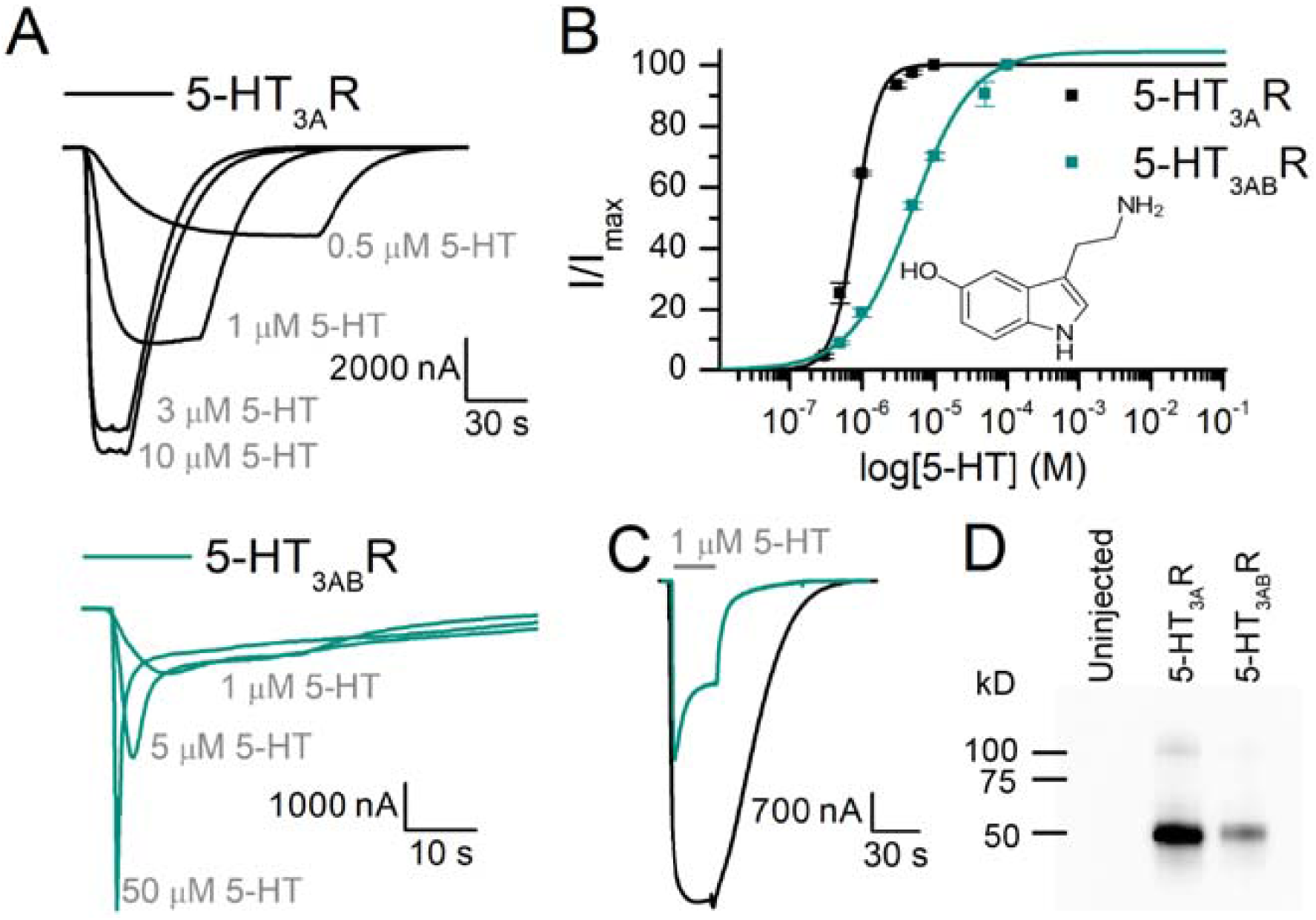
Comparing m5-HT_3A_R to m5-HT_3AB_R. (A) Sample traces of 5-HT_3A_R (black) and 5-HT_3AB_R (green) at varying concentrations of 5-HT. (B) Concentration-response curves show a higher potency for 5-HT with 5-HT_3A_R than 5-HT_3AB_R, as well as a steeper Hill slope. Parameters from these curves: 5-HT_3A_R: EC_50_ = 0.8 µM, n_H_ = 2.53±0.26, n = 5, 5-HT_3AB_R: EC_50_ = 4.30 µM, n_H_ = 1.04±0.02, n = 8. (C) Direct comparison of 5-HT_3A_R and 5-HT_3AB_R inward current evoked by 1 μM 5-HT for 30 s. (D) Western blot of membrane preparations of Xenopus oocytes expressing 5-HT_3A_R and 5-HT_3AB_R. V5-antibodies visualize 3A subunits containing V5-tag close to its N-terminus and highlight quantitative differences of the A subunit in 5-HT_3A_ and 5-HT_3AB_ receptors.

### Effect of Bupropion on 5-HT_3A_R and 5-HT_3AB_R

5-HT and a wide range of bupropion concentrations (A: 10 – 1000 µM; AB: 30 – 4000 µM) were co-applied to *Xenopus* oocytes expressing homomeric 5-HT_3A_R (Fig 2A, top) or heteromeric 5-HT_3AB_R (Fig 2A, bottom) under two-electrode voltage clamp. 5-HT was applied at a concentration that elicits approximately 30% of the maximal response (EC_30_, 5-HT_3A_R: 0.5 µM, 5-HT_3AB_R: 2 µM). Both traces in Fig. 2A show representative current responses at −60 mV. The first inward current represents the control current evoked by 5-HT alone, followed by subsequent currents obtained by the co-application of 5-HT (EC_30_) and increasing concentrations of bupropion, which dose-dependently inhibited 5-HT-induced currents for 5-HT_3A_ and 5-HT_3AB_ receptors. The concentrations inhibiting 50% of the control currents (IC_50_) were 87.1 µM (n_H_=1.28±0.07, n=5) and 866 µM (n_H_=2.07±0.20, n=7) for A and AB respectively (Fig. 2B; Table 1). Bupropion’s potency at 5-HT_3AB_R was 10-fold lower when compared to 5-HT_3A_R (unpaired t, t(10)=9.49, p=0.0000784). Notably, small potentiation of 5-HT-evoked currents was observed at low concentrations of bupropion (Fig. 2A bottom bar graph, 30 µM bupropion led to a 22.6% increase in current), and was therefore excluded from the IC_50_ fitting and analysis. Bupropion alone did not elicit an inward current for either receptor (Fig 2C).

**Figure 2.**
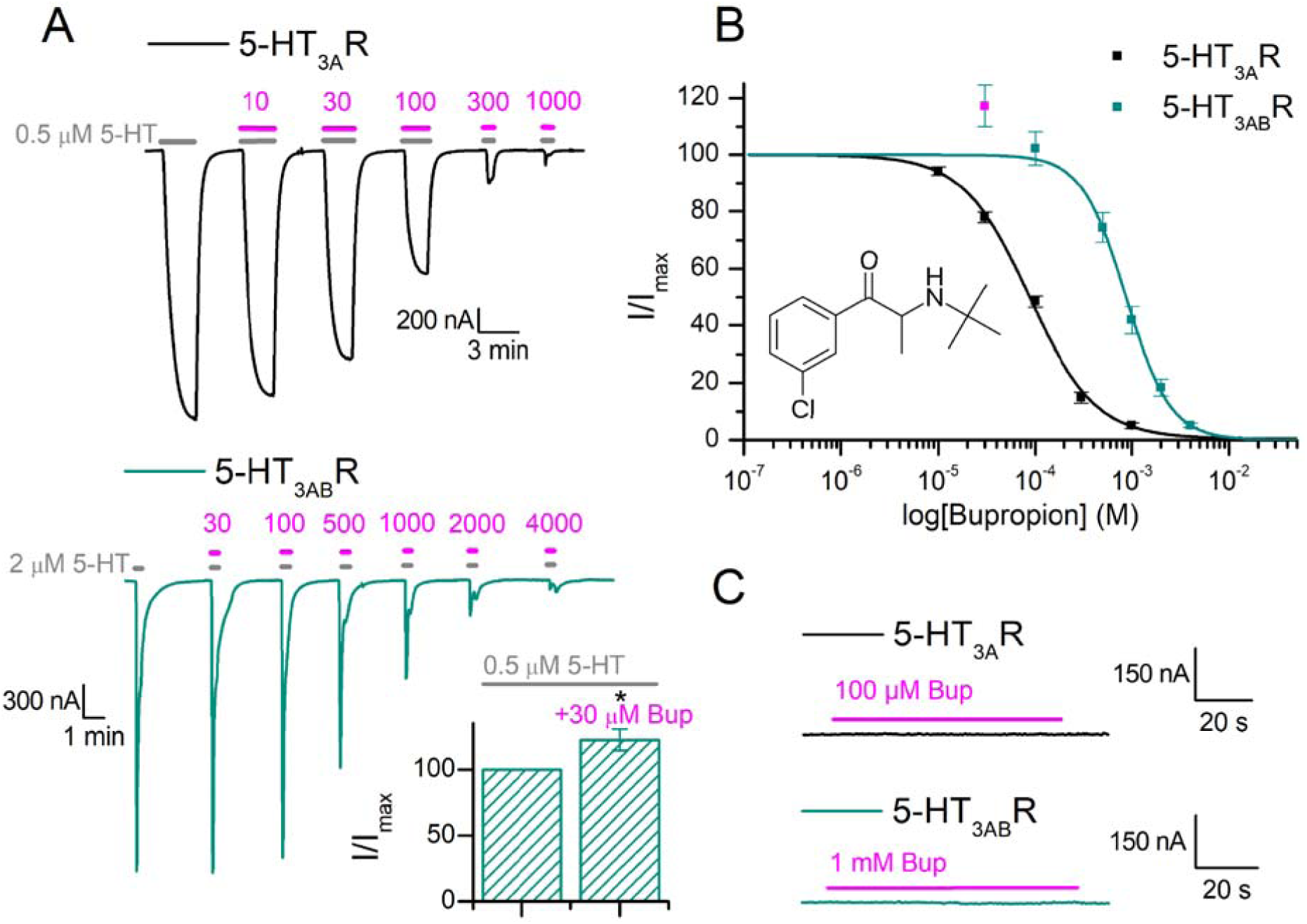
Bupropion’s antagonistic activity at homo- and heteromeric 5-HT_3_R. (A) Sample traces of oocytes expressing 5-HT_3A_R or 5-HT_3AB_R in response to 5-HT (∼EC_30_) alone and in combination with bupropion. 5-HT-evoked inward currents (grey, 5-HT_3A_ = 0.3 μM, 5-HT_3AB_ = 2 μM) were used as the control current. Following, the 5-HT concentration was kept constant and co-applied with increasing concentrations of bupropion (5-HT_3A_: 10 - 1000 μM, 5-HT_3AB_: 30 - 4000 μM). 5-HT_3AB_R bottom panel, bar graph showing that a low concentration of 30 µM bupropion in the presence of 2 µM 5-HT elicits potentiation (122.6±8.0%, n = 6) in 5-HT_3AB_R (B) Currents were normalized to the control currents and yielded the following IC_50_ values, 5-HT_3A_R: IC_50_ = 87.1 µM (n_H_ = 1.28±0.07, n = 5, means±S.E.M.), 5-HT_3AB_R: IC_50_ = 866 µM (n_H_ = 2.07±0.20, n = 7, means ± S.E.M.) (C) Oocytes expressing 5-HT_3A_R and 5-HT_3AB_R did not elicit an inward current in response to bupropion alone. Statistical significance was determined with paired t-test (*p ≤ 0.05 **p ≤ 0.01 ***p ≤ 0.001).

### Effect of Hydroxybupropion on 5-HT_3A_R and 5-HT_3AB_R

Hydroxybupropion, a major metabolite of bupropion, is involved in bupropion’s therapeutic effect, as it also inhibits DA/NE transporters, nAChRs, and 5-HT_3A_R (Bondarev, 2003; Damaj, 2004; Pandhare, 2017). Similar to bupropion, hydroxybupropion inhibited 5-HT_3A_R and 5-HT_3AB_R dose-dependently when co-applied with 5-HT (Fig 3A). The hydroxybupropion concentrations reducing the 5-HT-evoked currents to 50% of the initial response were 113 µM (n=5, n_H_=1.17±0.06) for 5-HT_3A_R and 505 µM (n=8, n_H_=1.57±0.12) for 5-HT_3AB_R (Fig. 3B; Table 1). Similar to bupropion, the potency of hydroxybupropion for 5-HT_3AB_R was right-shifted, resulting in a higher IC_50_ value when compared to 5-HT_3A_R, although to a lesser extent than bupropion (unpaired t, t(9)=16.26, p=0.000000185). Interestingly, low (50 μM) hydroxybupropion concentrations also led to small potentiation of 5-HT_3AB_R currents, which therefore were excluded from IC_50_ fitting and evaluations (Fig. 3A bar graph, average of 9.94% increase from initial current with 50 μM hydroxybupropion). Hydroxybupropion did not elicit a response in 5-HT_3A_ or 5-HT_3AB_ expressing oocytes when applied alone (Fig 3C).

**Figure 3.**
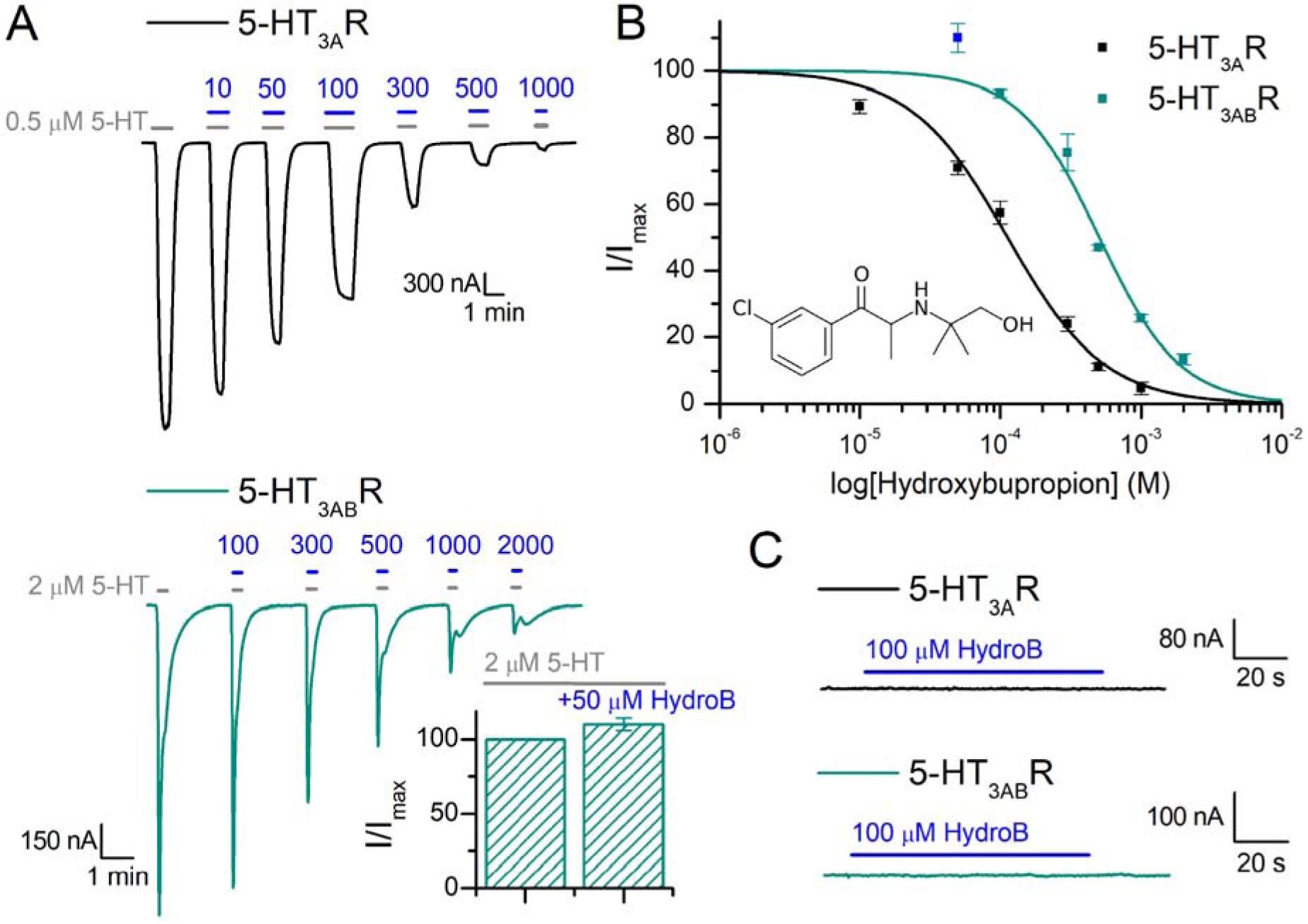
Hydroxybupropion is an antagonist for 5-HT_3A_R and 5-HT_3AB_R. (A) Sample traces of oocytes expressing 5-HT_3A_R or 5-HT_3AB_R in response to 5-HT (∼EC_30_) alone and in combination with hydroxybupropion. 5-HT-evoked inward currents (grey, 5-HT_3A_ = 0.3 μM, 5-HT_3AB_ = 2 μM) were used as the control current. Following, the 5-HT concentration was kept constant and co-applied with increasing concentrations of hydroxybupropion (5-HT_3A_: 10 - 1000 μM, 5-HT_3AB_: 50 - 2000 μM). 5-HT_3AB_R bottom panel, bar graph showing that a low concentration of 50 µM hydroxybupropion in the presence of 2 µM 5-HT elicits potentiation (109.9±4.31%, n = 5) in 5-HT_3AB_R (B) Currents were normalized to the control currents and yielded the following IC_50_ values, 5-HT_3A_R: IC_50_ = 113 µM (n_H_ = 1.17±0.06, n = 5, means ± S.E.M.), 5-HT_3AB_R: IC_50_ = 505 µM (n_H_ = 1.57±0.12, n = 8, means ± S.E.M.) (C) Oocytes expressing 5-HT_3A_R and 5-HT_3AB_R did not elicit an inward current in response to hydroxybupropion alone.

### Effect of Pre-incubation with Bupropion and Hydroxybupropion on 5-HT_3A_ and 5-HT_3AB_ Receptors

Bupropion’s allosteric inhibition of 5-HT_3A_R is not dependent on the opening of the receptor’s channel; it is non-use dependent (Pandhare, 2017). To evaluate the extent of inhibition evoked by pre-incubating oocytes expressing 5-HT_3A_ and 5-HT_3AB_R with bupropion or its metabolite, results were compared to the current amplitudes resulting from co-application of 5-HT and bupropion/hydroxybupropion. First, oocytes were perfused with 5-HT (∼EC_30_, 5-HT_3A_R: 0.5 μM, 5-HT_3AB_R: 2 μM) and bupropion (∼IC_50_, 5-HT_3A_R: 100 μM, 5-HT_3AB_R: 1 mM) to obtain the control current (Fig. 4A). Once a stable response was achieved, a constant IC_50_ concentration of bupropion was exposed to the receptors for exactly 5 min before another co-application of the same 5-HT and bupropion solutions. Pre-incubation decreased the current amplitude of 5-HT_3A_R to 76.1±3.20% (Fig. 4C, left panel) of control, consistent with previous findings (Pandhare, 2017). On the contrary, under the same experimental conditions, 5-HT_3AB_R was greatly affected by pre-incubation, which resulted in a current amplitude reduced to 35.5±2.29% of the control current (Fig. 4C, right panel). Similar results were obtained from hydroxybupropion pre-incubation (Fig. 4B). Compared to co-application alone, pre-application resulted in a greater depression of current for 5-HT_3A_R and 5-HT_3AB_R with hydroxybupropion (Fig. 4C, 5-HT_3A_R: 93.0±2.50% and 5-HT_3AB_R: 46.1±2.47% of control current).

**Figure 4.**
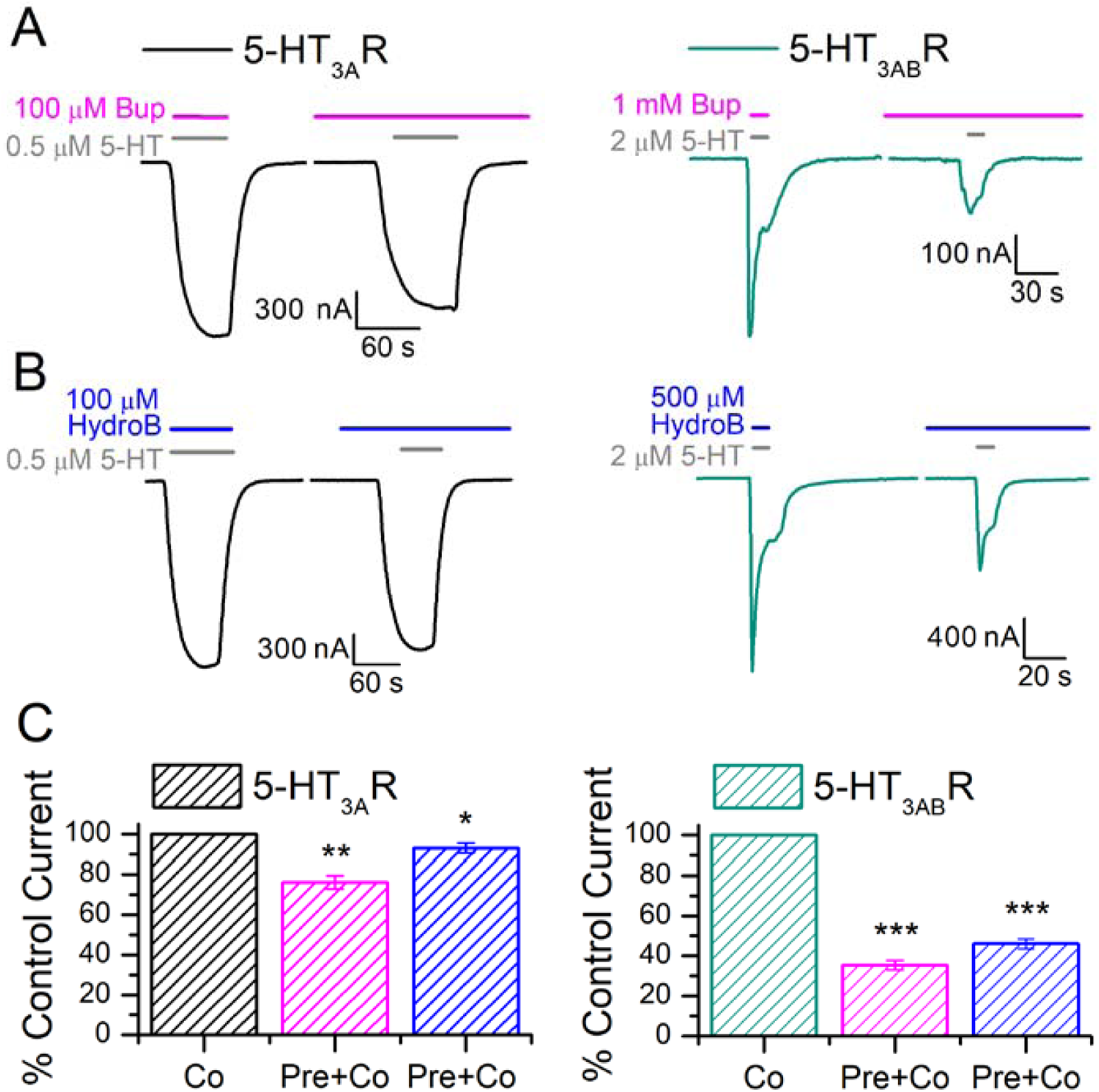
Non-use dependent allosteric inhibition of 5-HT_3A_R and 5-HT_3AB_R. (A) Sample traces of oocytes expressing 5-HT_3A_R (black, left panel) and 5-HT_3AB_R (green, right panel). The first 5-HT evoked currents were used as the control currents (grey bars, ∼EC_30_, 5-HT_3A_R: 0.5 µM, 5-HT_3AB_R: 2 µM) and was co-applied with bupropion (magenta bars, ∼IC_50_, 5-HT_3A_R: 100 µM, 5-HT_3AB_R: 1 mM). Following the stable 5-HT response, bupropion (∼EC_50_) was perfused for 5 min before another co-application of 5-HT and bupropion (B) Same experimental design as in (B), but with hydroxybupropion (blue bars, ∼IC_50_, 5-HT_3A_R: 100 µM, 5-HT_3AB_R: 500 µM). (C) Quantification of fractional inhibition of currents when the oocyte was pre-incubated in bupropion (magenta) or hydroxybupropion (blue) normalized to the control current (100%). Pre-incubation significantly reduced current amplitudes for 5-HT_3A_R (Bup: 76.1±3.20%, n = 5; HydroB: 93.0±2.50%, n = 6) and 5-HT_3AB_R (Bup: 35.5±2.29%, n = 6; HydroB: 46.1±2.47%, n = 4) as compared to co-application. Statistical significance was determined with paired t-test (*p ≤ 0.05 **p ≤ 0.01 ***p ≤ 0.001).

### Recovery times for Bupropion inhibition

Bupropion’s antagonistic effect on 5-HT-evoked inward currents has been shown to be reversible (Pandhare, 2017). To evaluate the recovery times for 5-HT_3A_R and 5-HT_3AB_R, bupropion at an IC_200_ concentration was applied for 60 s to show an observable effect on the oocyte’s current amplitudes and to better measure the time required to reverse the inhibitory effect. For these experiments, the ∼EC_50_ concentration of 5-HT (grey bars, 5-HT_3A_R: 0.8 μM, 5-HT_3AB_R: 5.0 μM) was applied episodically after washing in between each application (5-HT_3A_R: 2 min, 5-HT_3AB_R: 90 s minimum). These agonist-induced currents led to minimal run-down and the difference in current amplitudes was less than 5% (Fig. 5A). Sample traces of current recovery following bupropion application and removal are shown in Fig. 5B (left: black, 5-HT_3A_R, right: green, 5-HT_3AB_R). The first current response is the control current evoked by the agonist alone. Once a stable current response was obtained, bupropion was applied without the agonist for 60 s (magenta bars, ∼IC_200_, 5-HT_3A_R: 400 μM; 5-HT_3AB_R: 4 mM). Subsequently, the agonist was either applied immediately or after 10, 30, or 60 s (Fig 5B, top trace: 0 s, middle: 30 s, bottom: 60 s) after bupropion exposure. The largest decrease in current amplitude for 5-HT_3A_R was immediately after the bupropion application, leaving 82.4±1.78% of the initial current (Fig 5C, left panel). Increased wash times between bupropion and 5-HT applications led to a larger recovered current amplitude (Fig 5C, left panel). 5-HT_3AB_R showed a larger decrease in current amplitude after bupropion application and immediate 5-HT perfusion, not only with regard to the current amplitude but also its effect on the characteristically fast desensitization kinetics. Direct application of 5-HT after 60 s exposure to 4 mM bupropion led to a slow inward current that showed no desensitization but a slowly increasing current that never stabilized and was terminated after ∼3 min (Fig. 5B, first trace, right panel). After further washing, the fast activating inward current response exhibiting desensitization could be recovered. Due to the lack of a stable current response after 0 s of washing, the lowest current amplitude recorded was after 10 s of wash for 5-HT_3AB_R, with 18.1±2.19% of the initial current response (Fig 5C, right panel). Rapid recovery of current amplitude was achieved by increasing the wash times between bupropion and 5-HT applications (Fig 5C, right panel). 5-HT_3A_R and 5-HT_3AB_R both show stepwise and time-dependent recovery from bupropion inhibition, and can be fully reversed after 7+ min wash time.

**Figure 5.**
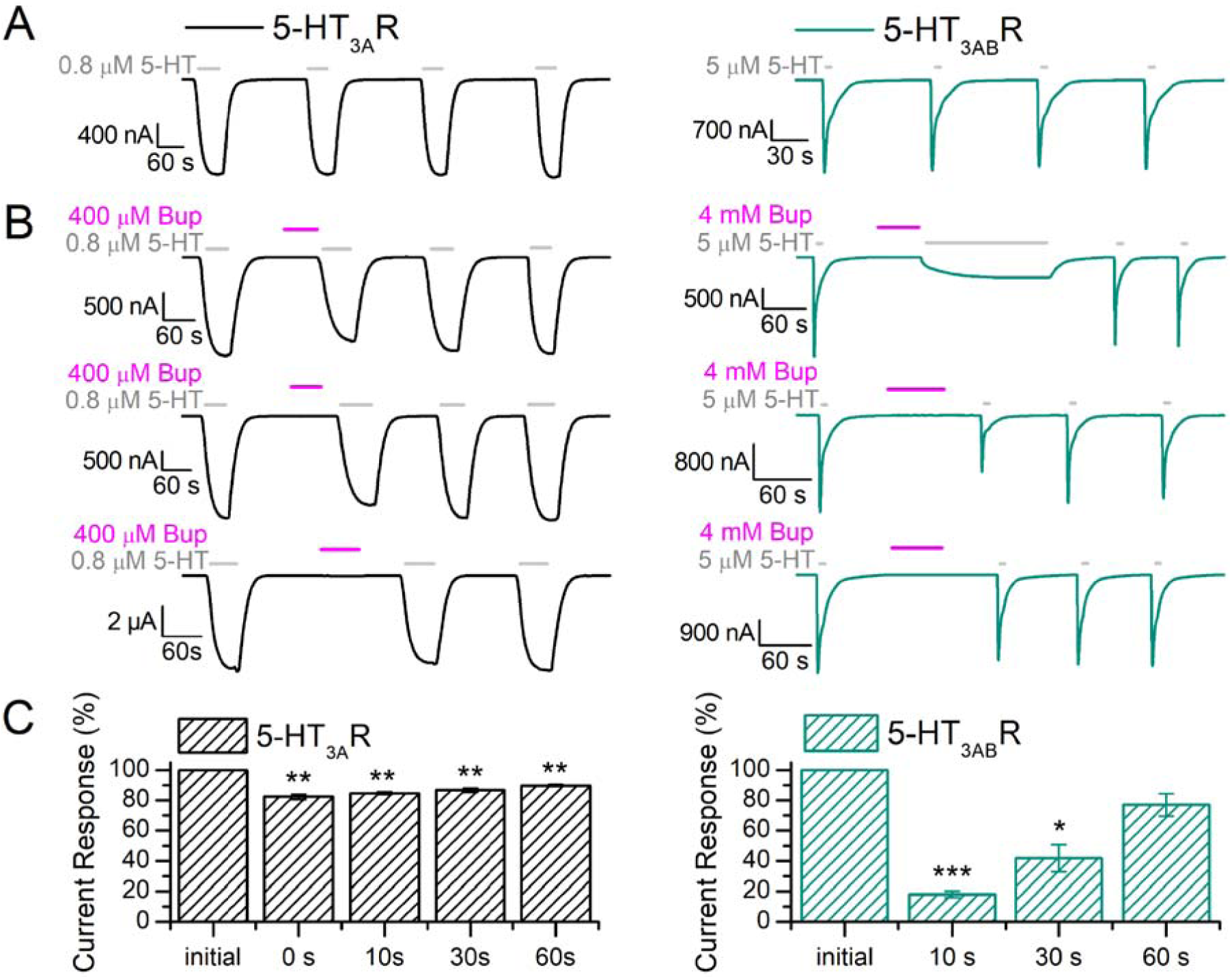
Recovery times for bupropion. (A)-(B) Shows sample traces of bupropion application (magenta bar) and the recovery times for 5-HT_3A_R (left panel, black) and 5-HT_3AB_R (right panel, green). (A) In two-electrode voltage clamp experiments, oocytes expressing 5-HT_3A_R and 5-HT_3AB_R showed a stable response to repeated applications of 0.8 μM and 5 μM 5-HT at −60 mV, with a minimal wash time of 2 min and 90 s between each application, for 5-HT_3A_R and 5-HT_3AB_R respectively. (B) The first 5-HT evoked response represents the control current for the recovery experiment. Bupropion (∼IC_200_, A: 400 μM, B: 4000 μM) was applied alone for 60 s at - 60 mV, followed by an immediate application of 5-HT. The grey and magenta bars represent the time of application of 5-HT and bupropion, respectively. Moving down the panel, the wash times after bupropion application were 0 s, 30 s, and 60 s. 5-HT_3AB_R current lacked desensitization and was slowly recovering for the 0 s wash after the application of bupropion and resulted in the application of 5 μM 5-HT for 3.5 min before withdrawing the agonist. (C) Quantitative representation of current amplitudes and results in (B) (n=3). 5-HT_3A_R was maximally reduced to 82.4±1.78% and 5-HT_3AB_R to 18.1±2.19% of the control current after 60s exposure to IC_200_ bupropion with 0 s and 10 s wash, followed by a stepwise recovery. All currents could be recovered to ∼95% after ∼7.5 min wash. Statistical significance was determined with paired t-test (*p ≤ 0.05 **p ≤ 0.01 ***p ≤ 0.001).

### Voltage-independent binding of Bupropion

To determine if bupropion binds to 5-HT_3_R in a voltage-dependent manner, 5-HT-induced currents (∼EC_50_; 5-HT_3A_R: 0.8 µM; 5-HT_3AB_R: 5.0 µM) were evoked in oocytes expressing 5-HT_3A_R and 5-HT_3AB_R at two different holding potentials, +40 and −40 mV (Fig 6A). First, the control current was obtained at positive and negative voltages before the co-application of 5-HT and bupropion (∼IC_50_; 5-HT_3A_R: 100 µM; 5-HT_3AB_R: 1mM). Bupropion reduced the current amplitudes of homo- and heteromeric receptors at both voltages. The mean fractional block was recorded at each voltage and normalized to the control current (Fig 6B; 5-HT_3A_R: 55.8±0.06%, 59.8±0.05%; 5-HT_3AB_R: 56.6±0.02%, 59±0.04%; −40 and +40 mV respectively, n=4). For 5-HT_3A_R and 5-HT_3AB_R, this fractional inhibition is similar at positive and negative voltages (paired t-test, 3A: t(3)=1.106, p=0.349; 3AB: t(3)=0.291, p=0.790). Based on these results, inhibition of 5-HT-induced currents by buproprion is independent of voltage.

**Figure 6.**
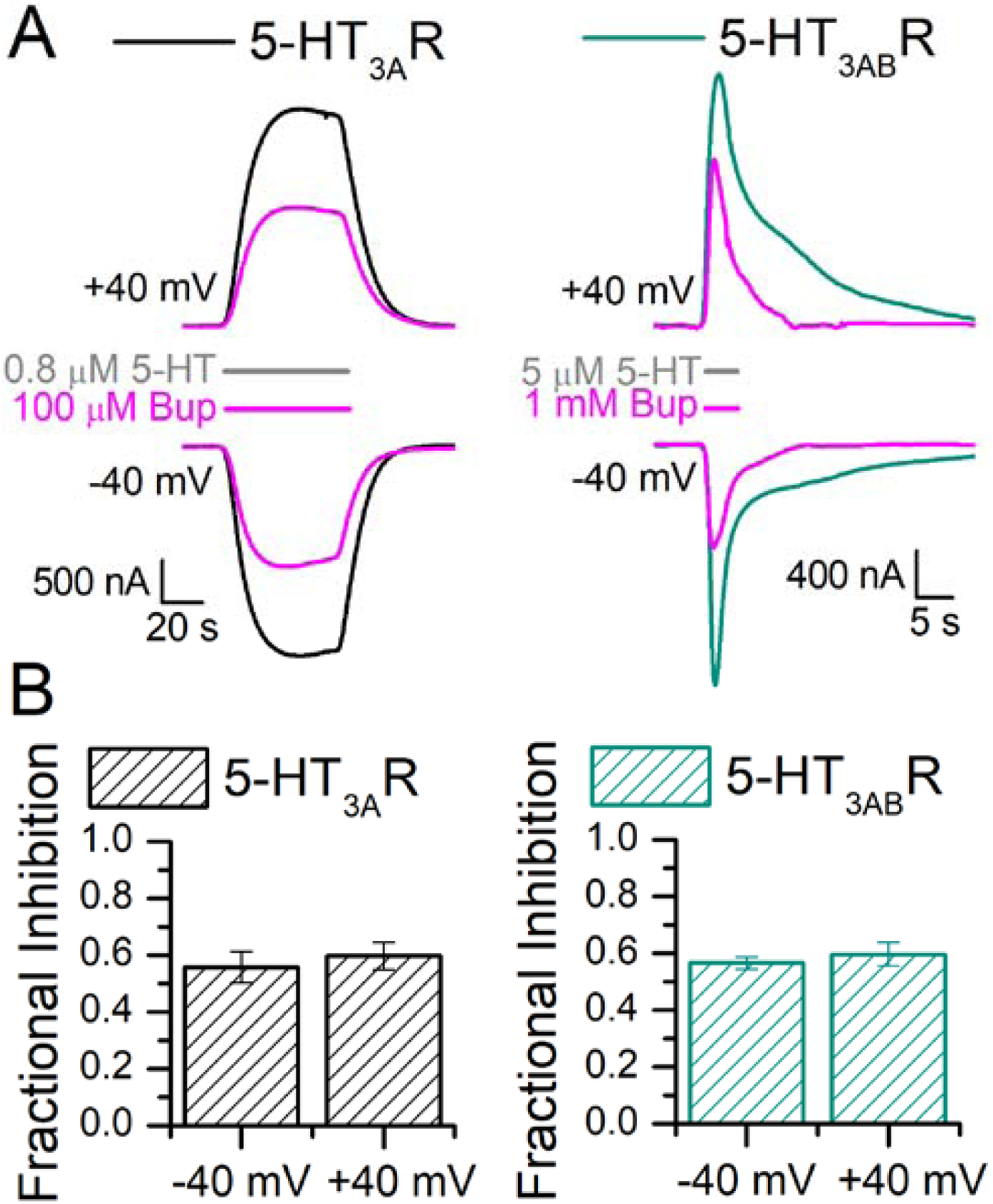
Voltage-independent block of 5-HT_3_-mediated currents by bupropion. (A) Sample traces of 5-HT_3A_R and 5-HT_3AB_R expressing oocytes (5-HT_3A_R: left, black; 5-HT_3AB_R: right, green) in response to 5-HT (∼EC_50_; top and bottom traces, 5-HT_3A_R: 0.8 µM; 5-HT_3AB_R: 5.0 µM) in the absence and presence of bupropion (magenta traces, ∼IC_50_; 5-HT_3A_R: 100 µM; 5-HT_3AB_R: 1mM) at different voltages. (B) Quantification of fractional inhibition, currents were normalized to the control currents at each voltage (n=4). Data is shown as mean±S.E.M. Statistical significance was determined with paired t-test (*p ≤ 0.05 **p ≤ 0.01 ***p ≤ 0.001).

### Bupropion at physiological concentrations and its effect on 5-HT_3A_R and 5-HT_3AB_R

To better understand the clinical significance of the bupropion-induced inhibition of 5-HT_3_R, 5-HT-induced currents were measured in the presence of a clinically relevant bupropion concentration (∼20 µM, (Schroeder, 1983; Vazquez-Gomez, 2014)). First, oocytes expressing 5-HT_3A_R and 5-HT_3AB_R were exposed to three different 5-HT concentrations (0.5, 1.0, 5.0 µM) in the absence of bupropion to obtain the initial current amplitudes (Fig 7A, black, left panel: 5-HT_3A_R, green, right panel: 5-HT_3AB_R). Next, the oocytes were continuously perfused with 20 µM bupropion and the same 5-HT concentrations were re-applied; the oocytes were pre-incubated with bupropion for at least for 2 minutes prior to 5-HT application (Fig 7A, magenta bars indicating bupropion presence). The results indicate that the continuous presence of a clinically relevant concentration of bupropion in the bath solution significantly inhibits 5-HT-induced currents of 5-HT_3A_R and 5-HT_3AB_R at all 5-HT concentrations tested (Fig 7B, paired t-test, p ≤ 0.05 or lower). Bupropion inhibited 5-HT-induced currents by ∼18% for 5-HT_3A_R (n=4), whereas 5-HT_3AB_R showed a ∼23% decrease in current (n=5, Fig 7B).

**Figure 7.**
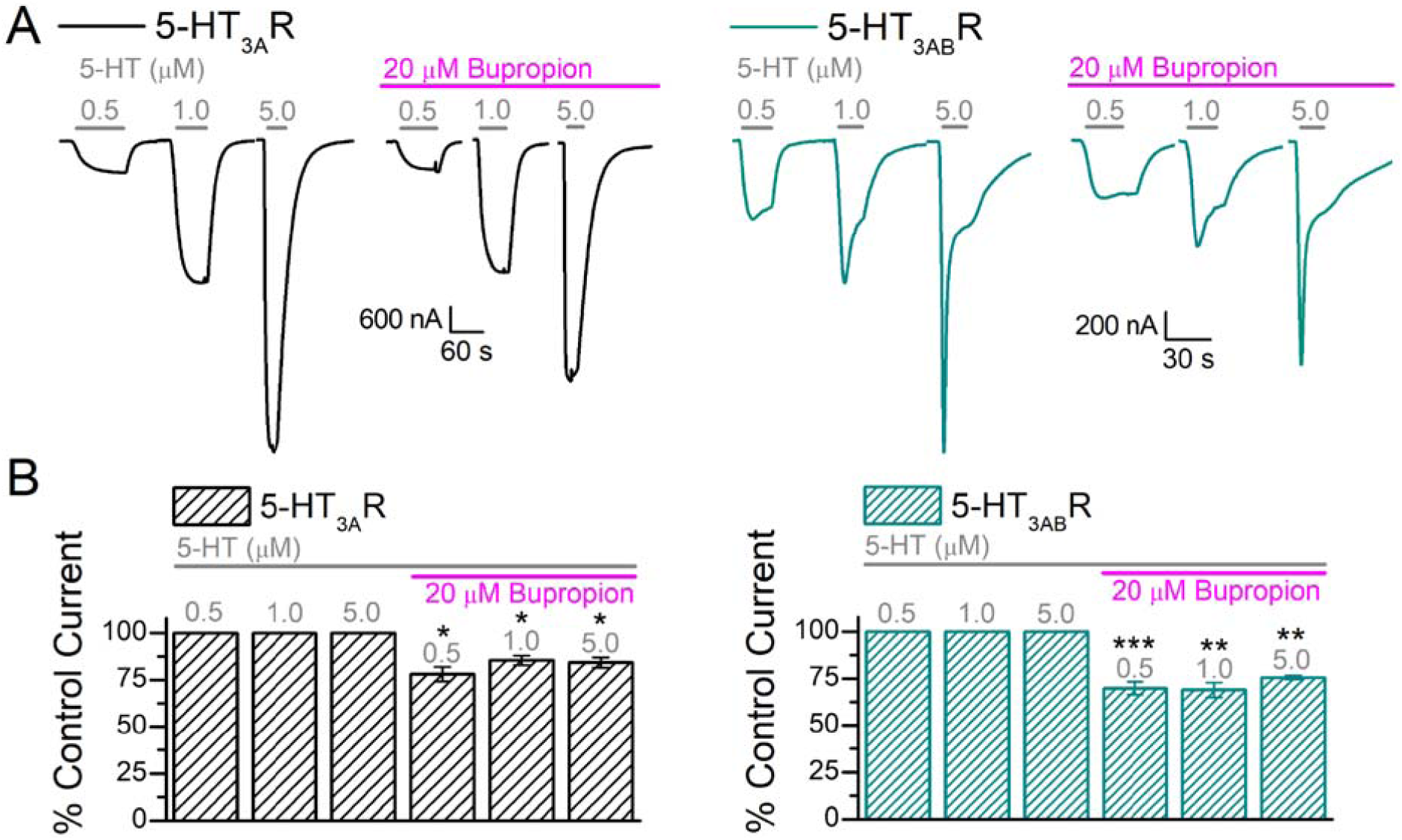
Bupropion at physiologically relevant concentrations and its effect on 5-HT_3A_R and 5-HT_3AB_R. (A) Sample traces of oocytes expressing 5-HT_3A_R (black, left panel) and 5-HT_3AB_R (green, right panel) in response to 0.5, 1.0, and 5.0 μM 5-HT (grey bars) followed by the same concentrations co-applied with 20 μM bupropion. Following the initial exposure to the three 5-HT concentrations (control current), the oocytes were exposed to 20 μM bupropion for at least 2 min before co-application with the agonist. (B) Quantitative representation of current amplitudes and results in (A) (A: n=4, AB: n=5). Data is shown as mean±S.E.M. Statistical significance was determined with paired t-test (*p ≤ 0.05 **p ≤ 0.01 ***p ≤ 0.001).

## Discussion

Our results, for the first time, demonstrate that bupropion antagonizes heteromeric 5-HT_3AB_ receptors and that the kinetics of inhibition are distinct from 5-HT_3A_R. Two-electrode voltage clamp experiments indicate that bupropion reversibly inhibits 5-HT-induced currents of *Xenopus* oocytes expressing 5-HT_3A_ and 5-HT_3AB_R in a concentration-dependent manner, with inhibitory potencies of 87.1 μM (same as previously reported (Pandhare, 2017)) and 866 μM respectively. Similar to other non-competitive antagonists (such as picrotoxin (Das, 2003)), bupropion has a lower potency (∼10-fold) at 5-HT_3AB_R compared to 5-HT_3A_ receptors and, therefore, could be used to discriminate between these two receptors (Thompson, 2013).

In humans, bupropion is extensively metabolized to the active metabolites hydroxybupropion and threohydrobupropion (Schroeder, 1983). One of these major metabolites, hydroxybupropion, is known to contribute to the biological efficacy of the parent drug (Bondarev, 2003; Damaj, 2004; Martin, 1990). Similarly to bupropion, the metabolite inhibits nAChR and 5-HT_3A_R in a non-competitive manner (Damaj, 2004; Pandhare, 2017) and also shares a comparable potency (unpaired t-test, p value=0.08464) for the homomeric receptor (5-HT_3A_R: IC_50_= 113 μM, similar to previously reported data (Pandhare, 2017)). In contrast, hydroxybupropion exhibits a ∼4.5-fold shift in IC_50_ (5-HT_3AB_R: IC_50_ = 505 μM).

Interestingly, bupropion and hydroxybupropion potentiate 5-HT-induced currents of 5-HT_3AB_R at low concentrations when co-applied with the agonist (Bup: 30 μM: 122.6±8.0%, n = 6; HydroB: 50 μM: 109.9±4.31%, n = 5), a phenomenon not observed with the 5-HT_3A_R. Similar potentiation has been observed for the antagonist atropine at neuronal nicotinic acetylcholine receptors (Parker, 2003; Zwart, 1997). Atropine potentiates acetylcholine-induced inward currents at low concentrations of the agonist but shows antagonistic activity at high concentrations of agonist or atropine. This phenomenon may be due to the drug acting on two distinct sites: one non-competitive inhibitory site and a second site associated with competitive potentiation (Zwart, 1997). Since we only observed potentiation in heteromeric 5-HT_3AB_ channels, this may indicate that the 3B subunits are mediating this effect. The 3B subunits potentially contribute directly to the potentiating binding site or mediate the effect via an allosteric mechanism. Bupropion-mediated inhibition of 5-HT_3A_R is non-use dependent (Pandhare 2017); the drug can access its binding site in a closed receptor state and inhibit the transition to an open state. In general, use-dependent block, or inhibition that would require a channel to be open to occur, is not influenced by pre-application. We evaluated the effect of a 5-min pre-incubation with bupropion and its metabolite hydroxybupropion on the homomeric and the heteromeric receptor (Fig. 4). Pre-incubation with inhibitor lead to an increased inhibition in all cases when compared to co-application, indicating that the block is non-use dependent for both receptors. Our observation that bupropion’s inhibition of 5-HT_3_R is voltage-independent additionally concurs with it not acting as an open channel blocker (Choi, 2003; Garcia-Colunga, 2011; Gumilar, 2003; Gumilar, 2008; Slemmer, 2000). Similar results are shown with other antidepressants at 5-HT_3_R (Eisensamer, 2003) and with bupropion at nAChR (e.g. α_3_β_2_, α_4_β_2_, α_3_β_4_) with predictions for an external binding site for bupropion (Garcia-Colunga, 2011; Slemmer, 2000). Considering different binding sites within the family of AChR (Pandhare, 2012), bupropion may have distinct binding sites in each channel (Arias, 2009b).

We saw a greater depression of current amplitudes when bupropion or its metabolite was pre-incubated as compared to co-application with 5-HT (Fig 5, 5-HT_3A_R: 76.2±3.20%, 93.0±2.50%; 5-HT_3AB_R: 35.5±2.29%, 46.1±2.47% of control current, Bup and HydroB respectively). During the pre-incubation experiments, bupropion binds and blocks the receptor prior to the opening of the channel, therefore presumably interacting with the closed channel and potentially inhibiting the channel from opening (Arias, 2009b; Choi, 2003). Consistent with other data, greater potencies of inhibition have been reported for bupropion and tricyclic antidepressants on Cys-loop receptors in the resting state than the open state (Arias, 2009b; Choi, 2003; Gumilar, 2008). This phenomenon may be due to the accumulation of antidepressants and antipsychotics in the cell membrane during pre-incubation, which may be important for the functional antagonistic effects of these drugs at the 5-HT_3_ receptor (Eisensamer, 2005).

Bupropion’s inhibition of 5-HT-mediated currents is reversible after a substantial amount of washing. In this study, we investigated the time it takes for 5-HT_3_R to recover from pre-incubation with bupropion at high concentrations (∼IC_200_; 5-HT_3A_R: 400 µM; 5-HT_3AB_R: 4 mM). Similarly to our pre-incubation experiments, bupropion reduced 5-HT_3A_R currents significantly but to a lesser extent than 5-HT_3AB_R. The largest reduction of current was observed with the shortest amount of wash time between the bupropion and agonist applications (5-HT_3A_R: 82.4±1.78% after 0 s wash; 5-HT_3AB_R: 18.1±2.19% of the control current after 10 s wash). For 5-HT_3AB_R, the immediate switch to 5-HT could not be quantified because the current never stabilized and was continuously increasing. This behavior is very uncharacteristic for 5-HT_3AB_R. We speculate that this high concentration of bupropion may accumulate in the membrane and stabilize the receptor in a closed position. In the 5-HT_3AB_R trace we may be observing the effect of bupropion being washed away over time during the application of the agonist, which results in the receptor opening and a very prolonged inward current. 5-HT_3A_ and 5-HT_3AB_ receptors show a time-dependent recovery from bupropion’s inhibition, and their currents could be fully recovered after ∼7.5 min of washing.

The clinical relevance of 5-HT_3_-inhibition by bupropion is currently unknown. Bupropion, but not its metabolites, concentrates in many tissues, with a brain to plasma ratio of 25:1 (Schroeder, 1983), which results in brain concentrations of ∼20 *µ*M (Vazquez-Gomez, 2014). Although 20 μM bupropion co-applied with agonist minimally inhibits 5-HT-induced currents of 5-HT_3A_R and possibly potentiates HT_3AB_R, pre-incubation with bupropion has a significant impact on its inhibitory effect (Fig. 7). Our results indicate that 5-HT_3_ receptors are significantly inhibited by 20 µM bupropion with a minimal pre-incubation time of 5 minutes (5-HT_3A_R: ∼82.7% 5-HT_3AB_R: ∼74.9% of control current). Moreover, hydroxybupropion also contributes to bupropion’s antidepressant activity (Bondarev, 2003; Damaj, 2004; Martin, 1990). The metabolite reaches ∼10-fold higher plasma concentrations in humans as compared to the parent drug (Findlay, 1981; Golden, 1988; Hsyu, 1997): with an average of almost 100 µM (based on clinical data, test ID: FBUMT; Mayo Clinic, MN), hydroxybupropion’s plasma concentrations are equivalent to 5-HT_3A_R’s IC_50_ value. Additionally, considering the increased inhibitory effect due to pre-incubation of 5-HT_3_R, we conclude that bupropion and hydroxybupropion have the potential to inhibit these receptors at physiologically relevant concentrations.

The comprehensive mechanism by which bupropion achieves its therapeutic efficacy is multifactorial. Bupropion inhibits nAChR and reuptake transporters for dopamine and norepinephrine. Due to low DAT occupancy in humans (∼26%: (Stahl, 2004)) it’s been hypothesized that dopamine reuptake inhibition is only partially responsible for its therapeutic effects in human subjects (Argyelan, 2005; Arias, 2009a). At therapeutic dosages, bupropion inhibits nACh receptors in the ventral tegmental area, dorsal raphe nucleus neurons, and interneurons in the hippocampal CA1 area (Alkondon, 2005; Arias, 2009a; Vazquez-Gomez, 2014). There, nAChR can modulate serotonergic projections (Aznar, 2005; Chang, 2011) and alter GABAergic transmission (Ji, 2000), in turn increasing dopamine levels, contributing to bupropion’s antidepressant activity (Arias, 2009a; Vazquez-Gomez, 2014).

5-HT_3_ receptors are localized in several areas involved in mood regulation, such as the hippocampus and prefrontal cortex (Bétry, 2011), and are found both pre- and post-synaptically where they modulate the release of many other neurotransmitters (Chameau, 2006; Faerber, 2007; Miquel, 2002; Nayak, 2000). For instance, 5-HT_3_ receptors and GABAergic neurons show strong interactions in the hippocampus and neocortical cells (Morales, 1996; Morales, 1997). Additionally, 5-HT_3_ receptors mediate stress-dependent activation of dopaminergic neurotransmission (Bhatt, 2013; Devadoss, 2010). Although 5-HT_3_ antagonist’s clinical relevance is mainly concerned with emesis, several studies have reported their potential role for treating mood and anxiety disorders (Bhatt, 2013; Thompson, 2007; Walstab, 2010). In humans, 5-HT_3_ antagonists show improvements of comorbid depression for patients suffering from complex disorders (Faris, 2006; Haus, 2000). Animal studies conclude that the antagonist’s anxiolytic activity is due to the inhibition of limbic hyperactivity responses (Bhatt, 2013), supported by the results that 5-HT_3A_R gene deletion produces an anxiolytic phenotype in mice (Kelley, 2003a). Furthermore, 5-HT_3_ antagonists have implications on hippocampal long-term potentiation (Bétry, 2011), increase synaptic NE levels, and facilitate 5-HT neurotransmission of other 5-HT receptors (Rajkumar, 2010). Also, several clinically available antidepressant and antipsychotic drugs have antagonistic activities at 5-HT_3_ receptors (Bétry, 2011; Choi, 2003; Eisensamer, 2003; Rammes, 2004). Interestingly, 5-HT_3_ antagonists have shown to enhance the antidepressant action of bupropion (Devadoss, 2010). In conclusion, 5-HT_3_ and nACh receptors have shown many implications in the neurobiology of depression and a highly complex interplay can be expected between these systems. Currently, it is not known if bupropion- or hydroxybupropion-mediated inhibition of 5-HT_3_ receptors is clinically relevant for their antidepressant activity. We show here that bupropion inhibits 5-HT_3_ receptors at clinically-relevant concentrations, and that this inhibition may contribute to bupropion’s desired and undesired clinical effects.

## Acknowledgments

Research reported in this publication was supported by the National Institute of Neurological Disorders and Stroke of the National Institutes of Health under award number R01NS077114 (to MJ). The content is solely the responsibility of the authors and does not necessarily represent the official views of the National Institutes of Health. We thank Dr. Myles Akabas (Einstein, NY) for providing us with the 5-HT_3B_ plasmid. We thank the TTUHSC Core Facilities: some of the images and or data were generated in the Image Analysis Core Facility & Molecular Biology Core Facility supported by TTUHSC.

## Author Contributions

A.G.S. and M.J. developed the study concept and design. A.G.S. performed experiments, acquired and analyzed data. A.G.S. and M.J. wrote the manuscript.

